# Reward expectation signals in the superior colliculus during associative learning

**DOI:** 10.64898/2026.09.11.750870

**Authors:** Sevinc Mutlu, Anindita Das, Marco Tripodi

## Abstract

Predicting future rewards allows neutral sensory cues to acquire motivational significance and guide goal-directed actions that maximise reward acquisition^1^. These reward predictions are updated when outcomes differ from expectations resulting in an *error* in prediction^2^. Midbrain dopamine neurons provide a canonical neural correlate of reward prediction error signal^3^ that are broadcast to multiple brain areas^4^. These learned reward predictions ultimately influence neural processes that enable animals to select, prepare and execute actions towards expected rewards^2,5^. Reward expectation signals have been observed across sensory and motor circuits where they can modulate sensory processing and action selection and vigor^6–8^. This raises the question of how integrative sensorimotor circuits represent information about reward expectation alongside sensory and motor variables that drive reward-seeking actions.

The superior colliculus (SC), a sensorimotor area in the midbrain traditionally associated with orienting and action selection^9–12^, is strongly influenced by reward context. Expected reward modulates visual and premotor activity, learned object values shape sensory responses, and recent reward alters visual processing in the mouse SC^13–16^. The SC also provides input to midbrain dopamine neurons and can relay or acquire conditioned sensory responses that influence dopaminergic activity^17–19^. However, SC projections can also drive orienting movements without supporting reinforcement^20^, raising the possibility that cue-related activity reflects salience or motor preparation rather than reward prediction itself. Here, using an associative-learning task in which auditory cues predicted distinct reward probabilities, together with longitudinal monitoring of intermediate SC population activity across learning, we asked whether the SC acquires signals related to reward probability, whether these signals can be dissociated from anticipatory licking, and whether outcome responses exhibit the hallmarks of reward-prediction errors. We found that cue-evoked SC activity progressively differentiated reward probabilities across learning, and that cue-associated reward probability explained neural activity beyond anticipatory licking. Thus, the intermediate SC carries a learned reward-expectation signal that is multiplexed with, but not reducible to, motor preparation. By contrast, outcome responses did not exhibit a consistent population-level signature of reward-prediction errors. Together, these findings identify the SC as a site where predicted reward is integrated with anticipatory action during associative learning.

## Results

Formal learning theories suggest that animals learn a causal association of cues with rewards when rewards are consistently associated with the cues in temporal proximity. This associative learning is predictive and leads to reward expectation^2,21^. Thus, reward expectation signals appear following conditioned cues representing a *prediction* of upcoming rewards, these responses are acquired during conditioning and exhibit amplitude modulation depending on probability of reward^22^. Additionally, canonical prediction error signals that enable associative learning is characterised by phasic peaks at reward delivery that scale with probability of reward and a phasic dip on unexpected reward omissions^3,22,23^. We asked if neural activity in the SC exhibits any of these features. Specifically, a) Does SC neural activity exhibit increased cue-evoked response during associative learning? b) Is the cue-evoked response modulated by cue-associated reward probability? c) Does SC neural activity signal a prediction error at outcome delivery?

To address these questions, we performed fiber photometric recordings of pan-neuronal calcium signals (AAV1-Syn-GCaMP6f) from the intermediate superior colliculus in head-fixed mice while training on a classical conditioning task. Mice were presented with auditory cues instructing 100% (100R), 20% (20R) or 0% (0R) reward probabilities and a fourth cue for 50% probability of an airpuff delivery (50P). We quantified anticipatory lick rate as the instantaneous lick rate within the window between cue onset and outcome (reward) delivery (Methods). For omission trials, the median outcome time of rewarded trials in that session was considered as the outcome time. Using this approach, we confirmed that reward-probability modulated anticipatory licking emerged during Pavlovian conditioning (Fig 1) with mean anticipatory lick rates in the last three sessions being overall higher for 100R trials than 20R trials.

**Fig 1:**
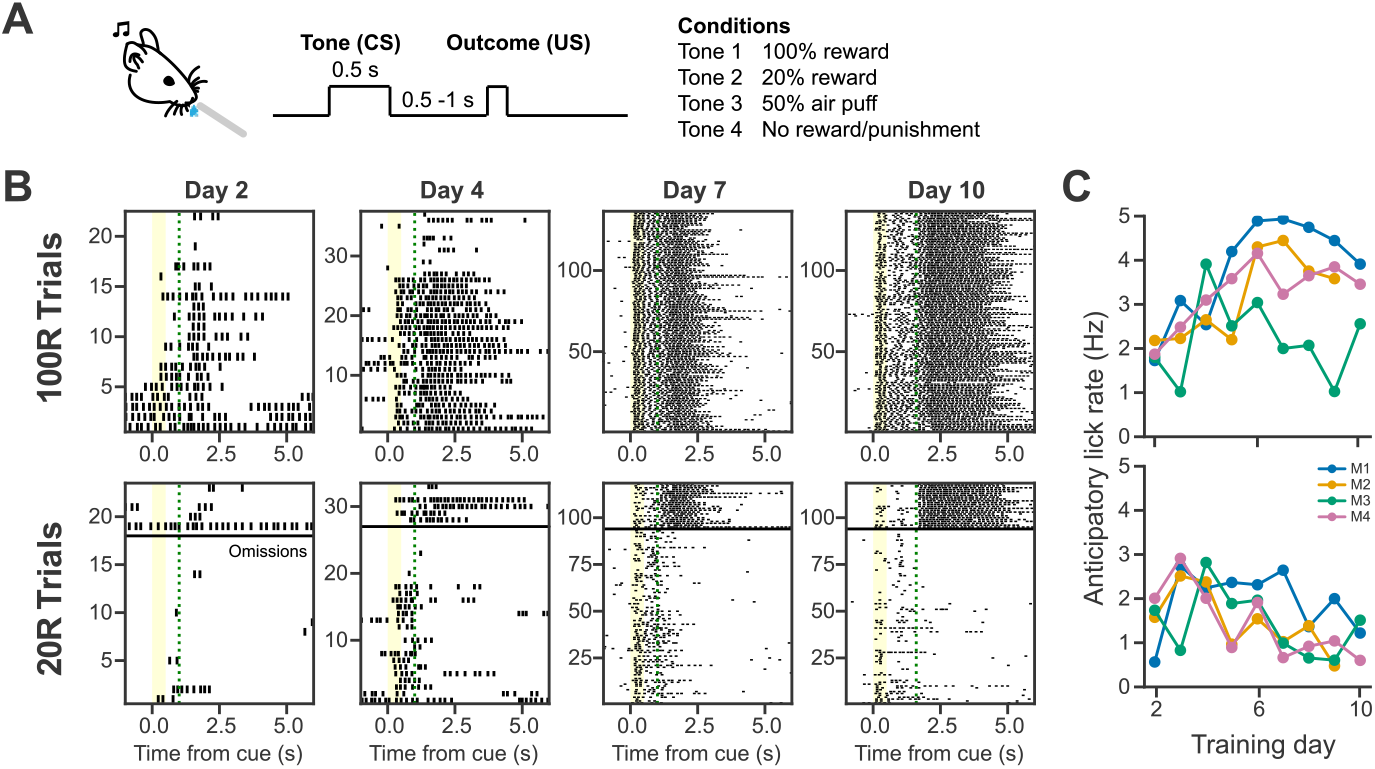
Reward modulated anticipatory licking during associative learning. **A)** Mice were trained on a classical Pavlovian conditioning task with auditory cues associated with different outcomes. *CS*: conditioned stimulus, *US*: unconditioned stimulus. **B)** Raster plots of licking for one example mouse show anticipatory licking following cue presentation (yellow bar) for trials associated with 100% (100R, top) and 20% reward probability (20R, bottom) respectively. For the 20R trials, omission trials are characterized by absence of stereotypical consumption licking following omissions (green dotted lines indicate time of reward delivery). **C)** Summary of mean anticipatory lick rates across sessions/days of training show higher overall lick rate at the end of training for the 100R trials compared to 20R trials.

### Reward expectation signals emerge in the superior colliculus during associative learning

Trial-aligned photometry traces showed a gradual change in dynamics over training days across mice (Fig 2A, Supplementary Fig 1A), with a strong rise following cue onset up until the peak following reward delivery and a gradual decay when rewards were withheld (orange dotted traces, Fig 2A). We henceforth refer to these as the pre-outcome and outcome epochs in the paper. Even though signal amplitudes and shapes, particularly post outcome, were variable across mice (Fig S1A), we noted a consistent pattern of response divergence between the two rewarded conditions during the pre-outcome epoch. To quantify the condition-wise signal modulation for the two epochs, we used the following metrics. For the pre-outcome epoch we considered the mean signal over the entire duration from cue onset up until outcome delivery. For the outcome epoch, we observed an increased signal amplitude during reward retrieval similar to previously reported studies^24,25^. To assess if this neural activity exhibited any prediction-error like modulation, we examined the rewarded and omitted cases separately. For rewarded trials, we quantified the phasic response by detecting the peak within 2s of reward delivery and then computed the mean signal amplitude within a small window (+/-200ms) around this peak. When rewards were withheld (omission trials), we noted that the signal did not exhibit a phasic dip. So, we measured the change in signal amplitude from the pre-outcome period – that is, the difference between the mean amplitude of the signal within the 2s outcome epoch and the signal amplitude immediately preceding (300ms) the outcome time. Similar analyses applied to the 0R and 50P trials (Fig S1B) showed little signal modulation. We therefore constrained our metric-based analysis of response modulation to the 100R and 20R trials to detect if signatures of reward expectation encoding were present in SC neural activity.

**Fig 2:**
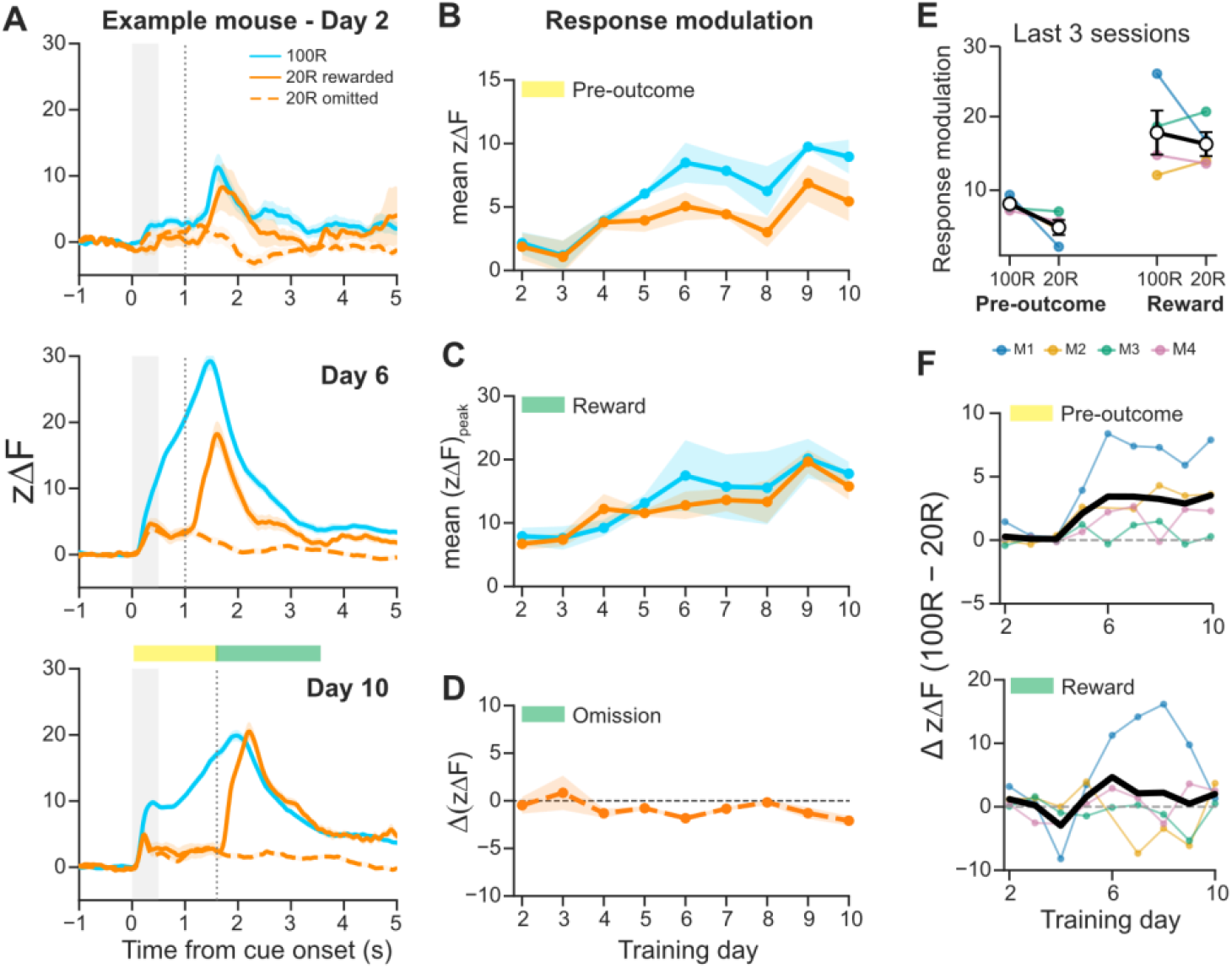
SC neural activity exhibit features of reward expectation signal. **A)** Trial-averaged calcium photometry traces for one example mouse during training. Yellow and green bars depict the pre-outcome and the outcome windows respectively used for quantifying epoch-wise response modulation metrics. **B)** Mean signal amplitude during the pre-outcome epoch were averaged across mice per day and plotted across training (shaded bars indicate standard error of mean) for the two reward conditions. The mean pre-outcome signal increased over learning with higher amplitudes for 100R trials than 20R. **C)** Phasic reward responses were quantified using peak detection over a window of 2s following reward delivery followed by the mean around this peak (+/-200ms). Mean phasic reward responses showed a progressive increase over training with no clear difference between the two rewarded conditions. **D)** Omission responses were computed by subtracting the mean amplitude of the signal over the 2s post-outcome duration from the mean signal preceding the outcome time. They showed very little change across training days. **E)** The mean response amplitudes towards the end of training are summarized for the two epochs for all mice individually and averaged across mice. **F)** A contrast metric (difference in response amplitudes between the two rewarded conditions for the pre-outcome and rewarded outcome epochs) confirmed the ‘directional’ influence of reward probability on anticipatory neural signal (F, top panel) across mice. This was notably absent for the phasic reward response (F, bottom panel) except for one mouse (blue trace).

We observed systematic changes in the pre-outcome signal with a progressive increase over training that differed by reward probability (Fig 2B, E). In contrast, while the phasic peak response to reward increased over learning they did not differ between the two conditions (Fig 2C, E). Statistical analyses (Methods) supported a significant change in both pre-outcome and phasic reward response across training days (*Response* ∼ day, Pre-outcome: *F*(8, 57) = 9.42, *p*_PERM_ = 0.0004; Reward: *F*(8, 57) = 6.60, *p*_PERM_ = 0.0026), however only the pre-outcome responses differed significantly between the two reward conditions (*Response* ∼ reward probability, Pre-outcome: *F*(1, 57) = 15.85, *p*_PERM_ = 0.0002; Reward: *F*(1, 57) = 1.18, *p*_PERM_ = 0.191). In contrast, during omission trials, SC signals primarily exhibited a decrease following withheld reward and this modulation didn’t change over learning (Fig 2D, *Response* ∼ day, *F*(8, 23) = 1.37, *p*_PERM_ = 0.262), suggesting that the change in SC neural activity during these trials may be consequent to either absent primary reward or cessation of licking (refer: raster plots for omitted trials in Fig. 1B bottom).

Our results demonstrate that the amplitudes of both the pre-outcome response and the phasic reward response evolve over learning. Significantly, the pre-outcome responses are modulated by the cue-associated reward probability, thus recapitulating a reward expectation signal. To assess if this modulation is observed consistently across mice, we computed a contrast metric by measuring the difference between the response metrics between the two reward conditions during the pre-outcome and the outcome epochs (Fig. 2F). In spite of the heterogeneity in the amplitudes of the responses as well as the temporal patterns of the learning trajectories across mice, the pre-outcome contrast metric showed a significant change over learning (*Contrast* ∼ day, Pre-outcome: *F*(8, 23) = 3.83, *p*_PERM_ = 0.0008; Reward: *F*(8, 23) = 0.55, *p*_PERM_ = 0.863).

Together, our results demonstrate that SC population activity exhibit features of reward expectation signals with reward probability modulated anticipatory neural response progressively emerging during associative learning (Fig. 2F). However, within the limitations of our current behavioural paradigm and recording methods, we did not observe features typical of reward prediction errors in the SC population activity.

### Neural-lick coupling is modulated by reward probability during associative learning

Given the primary sensorimotor function of SC and our observation that both anticipatory licking (Fig 1) and SC neural activity in the pre-outcome epoch (Fig 2) are modulated by reward probabilities, we wanted to decouple neural activity components related to reward probability from that associated purely with action execution. While this task did not involve head movements or saccades, studies have demonstrated the role of SC in goal-directed tongue movements^26–28^. We reasoned that if the changing SC activity predominantly reflected a change in ‘motoric’ drive, we would expect a) the neural signal to scale with licking throughout learning and b) the relationship between neural signal and licking to remain relatively stable across conditions and learning. Indeed, upon inspection of the trial-wise lick rates for individual mice across sessions, we observed that the SC activity in the pre-outcome epoch varied strongly with anticipatory licking (Fig 3A). To examine this relationship, we quantified the coupling between the neural signal and the licking behaviour using regression analysis. Given the distinct distribution of lick rates in the anticipatory and the reward delivery periods (compare Fig 3A with 3D top panel), we performed an epoch-wise regression analysis (Methods) for the two rewarded conditions (100R & 20R) separately:

**Fig 3:**
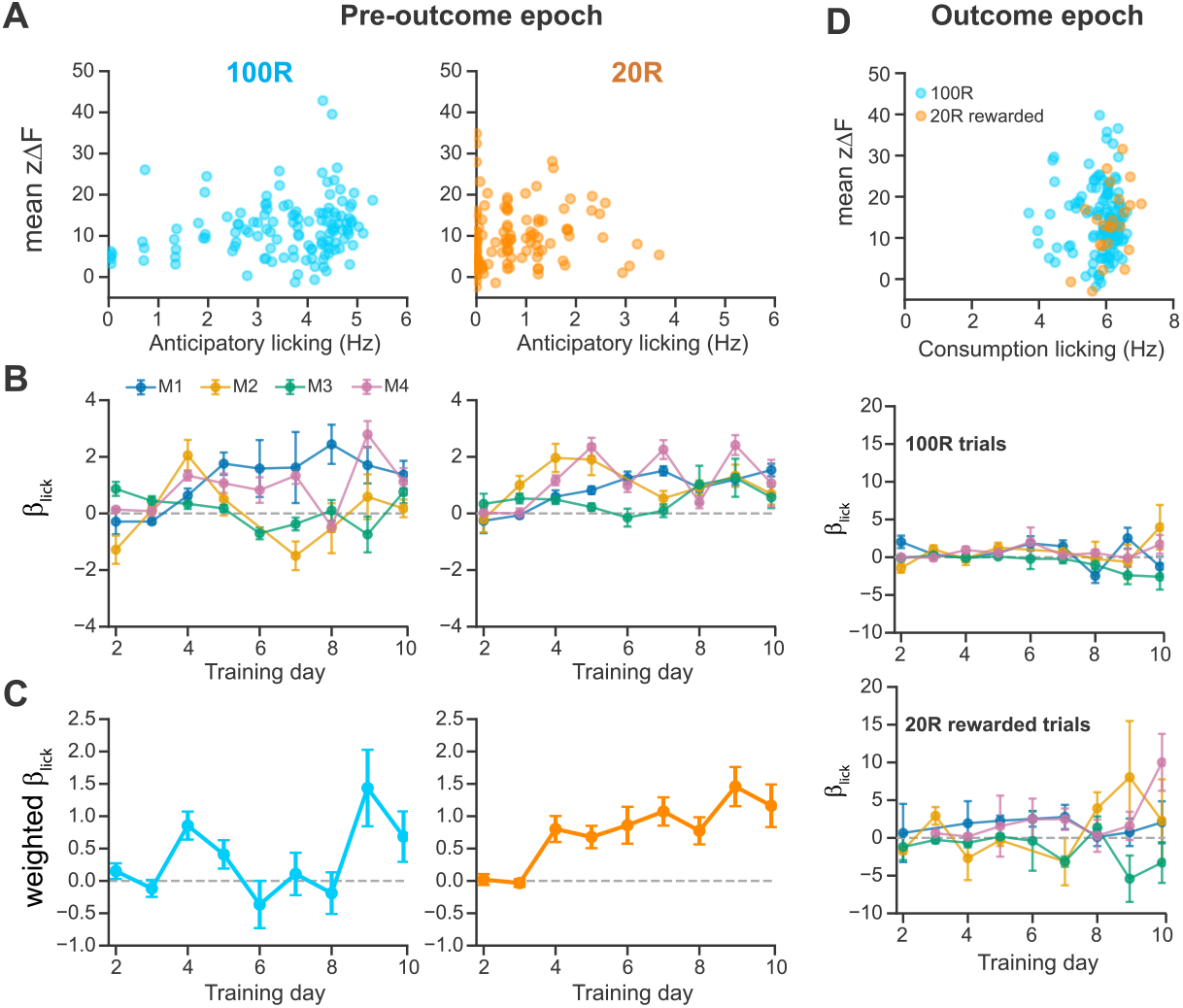
Anticipatory neural-lick coupling is modulated by reward probability. **A)** Scatter plots of the pre-outcome neural signal versus the anticipatory lick rate for one session from one example mouse show a correlation between the trial-to-trial fluctuations in neural signal and licking for the two rewarded conditions. **B)** Neural-lick coupling was quantified using regression analysis. The regression coefficient, *β*_*lick*_, was plotted across days for all mice individually and showed a gradual change in the neural-lick coupling with learning (error bars indicate standard error of mean). **C)** An inverse-weighted estimate (Methods) of the population *β*_*lick*_ showed a significant difference between the two rewarded conditions suggesting a reward probability modulated neural-lick coupling during the anticipatory phase. **D)** Similar analysis for the outcome epoch showed very little neural-lick coupling. Scatter plots showed an overall smaller dynamic range of trial-to-trial fluctuations in lick rates during reward retrieval. Regression analysis gave near-zero estimates of *β*_*lick*_ for both rewarded conditions.

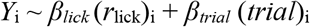

where, the regression coefficient, *β*_*lick*_, quantified the session-specific trial-by-trial coupling between neural activity (*Y*_i_) and lick rate (*r*_lick_) after accounting for the effect of trial position within the session (*trial*_i_). The term, *β*_*trial*_ (*trial*)_i_, accounted for the fact that both photometry signals and licking behaviour are subject to fluctuations over the course of a recording session, due to photobleaching effects or change in attentional or satiety state of individual mice respectively. The resulting session-level estimates for each mouse were then analysed across learning days to assess learning-related changes in the neural-lick coupling. Accounting for the variability across mice, we estimated a population neural-lick coupling trajectory by inverse-weighting the individual mice coefficients such that more precisely estimated coefficients (lower standard error of mean) contributed greater weight (Methods).

We observed that the neural-lick coupling within mice (Fig 3B) as well as their weighted population estimate (Fig 3C) became consistently positive later in learning for the probabilistic (20R) trials but fluctuated for the non-probabilistic (100R) case. While training day as a categorical factor did not significant improve the weighted linear model (Model *β*_*lick*_ ∼ day) for either reward conditions individually (100R: *F*(8, 23) = 2.13, *p*_PERM_ = 0.373; 20R: *F*(8, 23) = 7.38, *p*_PERM_ = 0.117), the learning-related trajectory differed significantly between the two reward conditions (Model *β*_*lick*_ ∼day*condition, *F*(8, 49) = 1.52, *p*_PERM_ = 0.005), implying a structuring of neural-lick coupling by reward expectation rather than a progressive increase in the coupling during learning.

Our results demonstrate that during associative learning, the SC neural dynamics following cue onset reflect the emergence of anticipatory licking that gets progressively shaped by learning of the cue contingencies. Interestingly, while mean anticipatory licking itself decreases for 20R trials over days (Fig 1) the mean neural signal amplitude during the pre-outcome period continues to increase (Fig 2) and the coupling between anticipatory licking and SC signal becomes strongly structured in the *uncertain* trials (Fig 3C). In other words, for probabilistic rewards, trial-to-trial fluctuations in SC activity following cue presentation is strongly coupled to the trial-to-trial variation in anticipatory action. In contrast, the strength of the coupling of the SC neural signal and consumption licking are much lower and doesn’t change with learning (Fig 3D). Together, our results suggest that while the learned cue-driven responses in the SC are expressed as anticipatory motor output, the SC population activity during associative learning reflects factors other than purely motor signals.

### SC neural activity exhibits multiplexed reward expectation and motor encoding

Our results thus far suggest that reward-expectation like neural dynamics appear in the SC during associative learning and this is expressed through contingent anticipatory behaviour. The regression analysis demonstrates that the coupling between neural activity and motor behaviour is particularly strong in the pre-outcome epoch compared to after reward delivery. Given these observations, we sought to answer how these different variables are encoded by SC neural activity during learning. In particular, we asked the following questions: a) How much of the neural activity is explained uniquely by the cue-associated reward probability after accounting for licking behaviour? b) Does the unique contribution of cue-associated reward probability and of licking change with learning?

Towards this end, we predicted the neural signal based on the task parameters and behavioural variables with an encoding model (Fig 4A, Methods). We considered a set of event predictors (cue, outcome), a cue-associated reward probability predictor and the behavioural variable (instantaneous lick rate). The event predictors (cue onset, rewards/omissions) were generated by convolving the respective binary predictor with a spline basis set to allow flexible temporal influence on the neural signal. A similar approach was used for the predictor representing reward probability, a probability-weighted temporally expanded predictor capturing the neural dynamics during the pre-outcome epoch, resembling a ‘reward expectation’ variable. For the full model, we tested kernels of different lengths and selected the one with the best performance based on all mice. We also confirmed that this final model improved predictions of the neural activity compared to ‘simpler’ models (Fig 4A bottom). We tuned this model based on the data from the last session of all mice and despite a variable model performance across mice, it captured the broad dynamics of the neural signals across days (Fig 4B).

**Fig 4.**
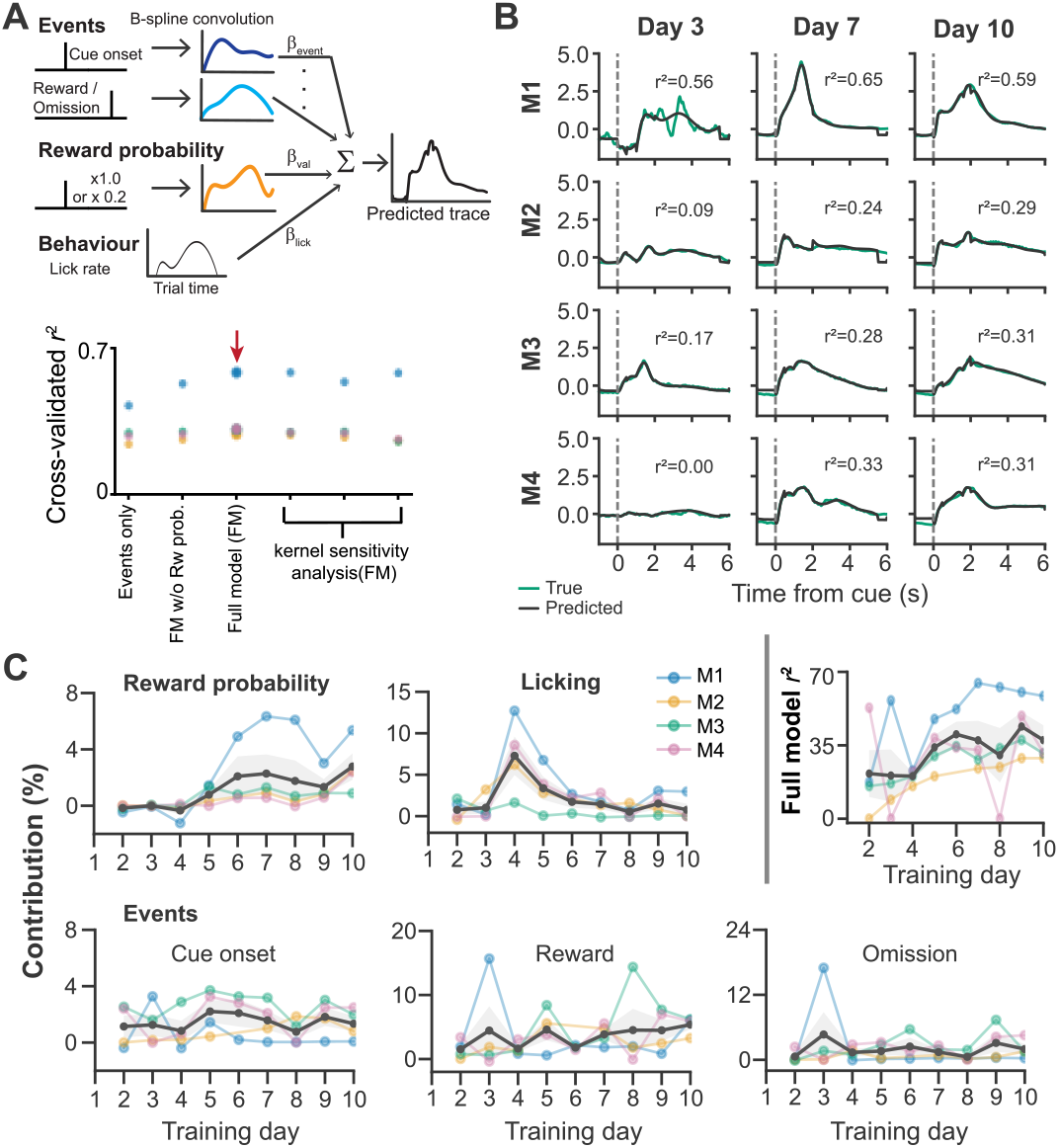
Multiplexed reward expectation and motor encoding in the SC. **A)** An encoding model with spline basis expanded event predictors, a lick rate predictor and a reward probability weighted temporally expanded cue predictor (cue valence) was used to predict session-wise neural traces. **B)** Predicted and actual traces for example sessions show that the model captured the broad neural dynamics across mice. **C)** The individual contribution of the behavioural and task-related variables to the neural signal were measured over learning. Cue valence contributed increasingly towards the neural signal over learning while lick rate contributed less during later sessions. There was no significant change in contribution of the event predictors across learning.

Using this encoding model, we quantified the unique contribution of each predictor to the neural activity by measuring the reduction in cross-validated predictive performance of the model when the predictor is removed (Methods). We performed this analysis for each session per mouse and plotted the change across days (Fig 4C). Across mice, the predictive performance of the full model improved during training confirming the increased representation of task- and behaviour-related variables in the SC neural activity over the course of learning. Despite the heterogenous range of values across mice, we observed key consistent patterns. Significantly, the unique contribution of cue-associated reward probability increased with learning. In contrast, the temporally expanded cue onset event without reward probability weighting did not show a significant change in contribution across learning (*F*(8, 23) = 0.84, *p*_PERM_ = 0.578). The unique contribution of lick rate showed a non-monotonic pattern, increasing initially and then decreasing for all the mice. While both cue-associated reward probability and licking showed a significant change in contribution across days (reward probability: *F*(8, 23) = 2.71, *p*_PERM_ = 0.0006; licking: *F*(8, 23) = 5.44, *p*_PERM_ = 0.0004), the event predictors did not (*p*_PERM_ > 0.05).

In addition, we examined the within-trial contribution of these predictors, focusing on only the reward-associated trials (100R & 20R). Using a functional regression analysis (Methods), we computed the time-resolved regression coefficients for individual predictors and assessed the change in coefficient curves across learning. This showed a largely heterogeneous pattern across mice with licking and reward probability contributing to the pre-outcome activity increasingly over learning, but their exact temporal patterns varied across individuals (Fig S2).

In conclusion, our results suggest a multiplexed encoding of cue-associated value and action within the SC resembling primary somatosensory and motor cortical activity in primates^8^. Importantly, this offers a possible mechanism by which value-guided action selection could occur at the level of the SC either by directly influencing actions controlled by SC such as orienting responses or by sending this information to downstream targets such as the dopaminergic-striatal circuit. Future studies could probe this using more complex and natural behaviours involving SC-directed orienting actions. This along with neural recordings including cell-type specific recordings could further dissociate if the heterogeneity in our results reflects different neuronal types contributing to value and motor encoding^29^ or a convergence of inputs onto single SC neurons from value-encoding circuits and motor circuits controlling anticipatory actions^30^ or other latent behavioural variables not directly measurable in this study.

## Discussion

Our results demonstrate that population activity in the intermediate superior colliculus carries a reward-expectation signal that emerges during associative learning. Following cue presentation, anticipatory SC activity progressively differentiated cues predicting high and low reward probability, showing that the developing response reflected the learned reward contingency rather than the sensory cue alone. Critically, this modulation could be dissociated from purely motor preparatory activity. During 20% reward trials, anticipatory licking decreased across learning while the mean pre-outcome SC response continued to increase, demonstrating that neural activity did not simply scale with the magnitude of the forthcoming action. Moreover, cue-associated reward probability explained unique neural variance after instantaneous licking had been accounted for, and its contribution increased over learning as the contribution of licking declined. Together, these analyses establish that the SC represents reward expectation independently of, but alongside, the anticipatory actions through which that expectation is expressed.

This finding extends the established role of the SC beyond sensory processing and movement generation. While it has been previously demonstrated that neurons in the SC exhibit *predictive* activity prior to generating saccades^31^ which can causally mediate target selection in primates^32^ and mice^33^, our results suggest that the SC contains prospective reward-expectation signals even in the absence of visual targets and when the animals are not required to perform saccades complementing studies supporting a more general role of the SC in cue-guided action selection^34,35^. Moreover, reward expectation is known to modulate visual and premotor responses in the primate SC, while recent reward history shapes visual processing in the mouse SC^13–16^. Our results show that reward-related modulation is not simply imposed on an otherwise motor signal. Instead, reward probability becomes an explicit component of SC population activity through learning. The SC therefore appears to integrate predicted value with action-related signals, placing it in a position to bias the selection and preparation of actions according to their anticipated outcomes. Such signals could influence behaviours controlled directly by the SC^10,36–38^ or be transmitted through collicular projections to dopaminergic and striatal circuits ^17,18,39^ involved in reinforcement learning and value-guided action selection^5,40^.. The strong contingency-dependent coupling between SC activity and anticipatory licking may therefore reflect the conversion of reward expectation into an appropriate behavioural response, rather than a motor confound.

By contrast, the data do not collectively support a robust, canonical reward-prediction-error signal at the level of SC population activity. Classical prediction errors reflect the difference between expected and obtained outcomes, producing larger responses to unexpectedly delivered rewards and negative responses when expected rewards are omitted^3,22^. In our recordings, phasic reward responses changed across learning but were not systematically larger when reward followed the 20% cue than when it followed the fully predictive cue. Negative-going responses following reward omission were observed on some individual training days, but they were variable in timing and magnitude and were diluted when responses were aggregated across learning. These occasional responses may reflect transient prediction-error signals at particular stages of learning or activity confined to subsets of SC neurons, but they do not constitute a stable population-level signature. A limitation of the present study is that outcome-related prediction-error signals were assessed using only 100% and 20% reward contingencies and through pan-neuronal fibre-photometry recordings. Although these conditions provide a substantial difference in reward expectation, they do not establish whether outcome responses scale systematically across a broader range of predicted reward probabilities. Moreover, omissions occurred only following the 20% cue and were therefore relatively expected, limiting the ability of the task to reveal robust negative prediction errors. Fibre photometry may further reduce sensitivity to such signals by averaging activity across heterogeneous neuronal populations. Prediction errors carried by sparse neuronal subsets, expressed with opposite signs across cells, or occurring with variable timing could therefore be diluted in the population signal. Future studies should combine more finely graded reward contingencies with rare unexpected reward and omission probes, and use electrophysiology or cell-resolved calcium imaging to determine whether individual SC neurons encode positive or negative prediction errors and how these responses combine at the population level. Such experiments will determine whether the lack of a consistent prediction-error signature in bulk SC activity reflects the absence of such signals in the SC or their dilution through the averaging of temporally and functionally heterogeneous single-neuron responses.

Overall, our findings establish that the intermediate SC carries a learned reward-expectation signal that is separable from pure motor preparation. Rather than representing reward expectation and action in isolation, SC population activity multiplexes these variables, providing a potential substrate through which predicted value can directly shape action selection.

## METHODS

### Animals & Surgery

Four adult C57BL/6 wild-type mice were used for this study. All animal procedures were conducted in accordance with the UK Animals (Scientific procedures) Act 1986 and European Community Council Directive on Animal Care under project license PPL PCDD85C8A and approved by The Animal Welfare and Ethical Review Body (AWERB) committee of the MRC Laboratory of Molecular Biology.

Mice were anaesthetized with isoflurane. Viral injections in the brain were done using a nanoject (Scientific Laboratory Supplies) equipped with a pulled borosilicate glass capillary. A maximum of 300 nl of virus (AAV1-Syn-GCaMP6f) was injected within the lateral SC along with fibre implanted just dorsally to the intermediate SC. Post-surgery, mice were housed individually to prevent damage to implants. Lighting was set to a reversed light:dark cycle, with simulated dawn and dusk at 19:00 and 07:00, respectively. Temperature was maintained at 19-23 °C and humidity at 45–65%. Mice were food restricted to maintain 85% of their free-feeding weight.

### Behaviour & video data processing

Mice were trained on a head-fixed setup with auditory cues delivered through speakers instructing the different contingencies – 100% reward, 20% reward, 0% reward and 50% punishment (air puff delivery). Behavioural events were controlled and recorded by Unity-based task software with video recordings of the animal throughout each recording session. Task-event timestamps, including auditory cue onset and outcome delivery, were recorded by Unity and used to align the photometry signal to individual trials. On Day 1 of training, mice were habituated to head-fixing followed by introduction to the baseline task (cue followed by reward 100% of times) to encourage licking. The probabilistic reward condition (20%) was introduced only on the second day. The no-reward and punishment trials were introduced from day 4 and day 5 onwards. All mice were recorded for a total of ten days.

#### Video-based behavioural analysis and alignment

Behavioural videos were acquired at 40 frames/s and analysed using DeepLabCut (DLC)^41^. DLC tracking included the tongue and eye, together with an LED visible in the behavioural video that provided a trial-timing marker. Trial onset in the video was identified from the DLC likelihood trace of the LED using threshold-based detection of transitions into the LED-on state, with detected onsets required to persist for at least two consecutive frames. Video-frame indices were converted to time using the 40-Hz camera frame rate, and licking and blinking events detected from the DLC likelihood traces were expressed relative to the corresponding video-derived trial onset.

Licks were detected from peaks in the DLC tongue-likelihood trace using a minimum peak prominence of 0.24, likelihood threshold of 0.75 and minimum separation of two video frames. Blinks were identified from peaks in the inverse eye-open likelihood trace, using a minimum prominence of 0.55 and minimum separation of 120 frames. Detected LED, lick and blink frames were converted to time using the video acquisition rate of 40 frames/s. Each lick and blink event was then expressed relative to the corresponding LED-defined trial onset, and events occurring from 3 s before to 10 s after trial onset were retained for each trial.

DLC-derived trials were matched sequentially to the corresponding trial-wise photometry/Unity dataset after removal of the first video trial, consistent with the first-trial exclusion applied during photometry preprocessing. Trial counts were required to match before the datasets were merged. The correspondence between video-derived and Unity/photometry trials was verified by comparing trial counts and inter-trial intervals across the two data streams before merging the datasets on a trial-by-trial basis. Following verification of alignment, DLC-derived behavioural events were combined with the corresponding Unity trial information and trial-aligned photometry traces for subsequent analyses.

### Neural recordings & photometry data pre-processing

Fibre photometry recordings were acquired using a Doric photometry system. GCaMP and isosbestic reference fluorescence signals were recorded simultaneously and demodulated during acquisition. Signals were sampled at 120 Hz, processed and synchronized with behavioural task events for subsequent trial-aligned analysis. Photometry signals were processed separately for each recording session. Raw GCaMP and isosbestic reference signals were visually inspected for large transient artifacts. Where such artifacts were present, affected samples were identified using a robust median absolute deviation-based threshold and replaced by linear interpolation before further preprocessing. The resulting signals were smoothed using a centred 10-sample moving average. Slow baseline drift in each channel was estimated using adaptive iteratively reweighted penalized least-squares (airPLS) fitting and subtracted from the corresponding smoothed signal. The baseline-corrected calcium-dependent and reference signals were then independently z-scored across the recording session. To correct for fluctuations shared between the two channels, the z-scored isosbestic reference signal was fitted to the z-scored calcium-dependent signal using robust linear regression with Tukey bisquare weighting. The fitted reference signal was subsequently subtracted from the calcium-dependent signal to obtain the z-scored, reference-corrected fluorescence signal, defined as:

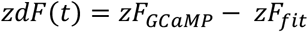

where, *zF*_*fit*_ = *β*_0_ + *β*_1_*zF*_*ref*_ obtained from the robust linear fit of the z-scored reference (isosbestic) signal to the z-scored GCaMP signal. The resulting corrected and standardised photometry signal, *zdF*(*t*) was aligned to auditory cue onsets per trial to obtain cue-aligned traces for subsequent analysis.

### Behaviour and neural metrics computation

The primary measurement of behavioural performance in our task is licking. Instantaneous lick rates were quantified on a trial-by-trial basis based on trial-aligned lick events detected from the video analysis. Binned lick events were smoothed with a gausian kernel of 150ms to obtain the trial-aligned lick rate trace. Anticipatory licking was calculated during the interval preceding outcome delivery, while consummatory licking was quantified following reward delivery. Actual reward delivery times for each trial were used, whereas, for omission trials, the median outcome time of rewarded trials in that session was considered the outcome time.

Summary neural activity metrics were computed for the pre-outcome and post-outcome epochs respectively. For the pre-outcome epoch the mean signal over the entire duration from cue onset up until outcome delivery was considered as the measure of the anticipatory neural activity. For the outcome epoch, rewarded and omission trials were treated separately. For rewarded trials, the phasic reward response was quantified by detecting the peak within 2s of the reward delivery and then computing the mean within a small window (+/-200ms) around this peak. For omission trials, the change in signal amplitude was measured by computing the difference between the mean amplitude of the signal within the 2s ‘reward’ epoch and the mean amplitude immediately preceding (300ms) the outcome time.

All metrics were first calculated for individual trials and subsequently summarised within each recording session and trial condition. Training sessions were assigned to their corresponding experimental training day, preserving gaps for days on which no recording was available due to corrupt neural or video data. Based on this only one day was missing for one mouse (total = 35 sessions across mice). Since the probabilistic reward condition was only introduced on second day of training, we began our learning-related analysis from Day 2 and have identified it as such in all figures. For analyses examining learning-related changes, we compared session-level measures across training days while treating individual animals as the biological replicates.

### Regression analysis

We quantified the relationship between trial-by-trial licking behaviour and photometry responses using linear regression. This neural-lick coupling was quantified separately for the pre-outcome and outcome epochs as defined above. For the pre-outcome analysis, the neural response metric described above was used, whereas outcome responses were quantified simply as the mean z-scored fluorescence during the 2 s following the trial-specific outcome time. Outcome analyses included all 100R trials and rewarded 20R trials only. For each mouse, training day and reward condition separately, trial-wise neural responses were modelled using ordinary least-squares regression as follows:

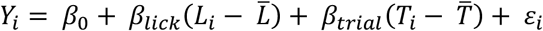

Where *Y*_i_ is the neural response on trial *i, L*_i_ is the corresponding lick rate and *T*_i_ is the trial number within session. Both lick rate and trial number were mean-centred for a given condition within a session for every mouse. Thus, *β* _lick_ quantified the trial-by-trial relationship between licking and neural activity while accounting for systematic changes in neural responses over the course of the recording session. We use this regression coefficient as a measure of the neural-lick coupling, with its standard error providing an estimate of the uncertainty of the session-level coefficient estimate. For the pre-outcome epoch, it represents the coupling between neural activity and anticipatory licking, while for the outcome epoch it quantifies the coupling with reward consumption licking.

Finally, for visualization of the population trajectory, session-level estimates were combined across mice using inverse-variance weighting,

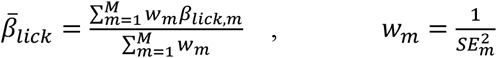

such that more precisely estimated coefficients contributed more strongly to the population estimate, 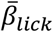 with the standard error of the population estimate 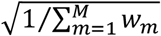

Statistical inference on learning- and condition-dependent changes was performed separately using permutation tests.

### Encoding model

Photometry signals were modelled using a linear encoding framework as follows

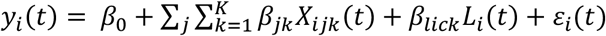

Where *y*_*i*_(*t*) is the photometry signal on trial *i* at time *t*; *X*_*ijk*_(*t*) represents the *k*-th temporally expanded event predictor *j, β*_*jk*_ is its fitted component, *L*_*i*_(*t*) is the continuous lick rate predictor and *β*_*lick*_ its fitted component. Neural activity and predictor columns were z-scored across all time points before model fitting.

The event predictors comprised cue onset, reward delivery and reward omission. For the trials associated with 0% reward and the punishment trials, the outcome predictor was assigned as a reward omission. To explicitly assess contribution of an *expected reward* we used a temporally expanded cue predictor weighted by its associated nominal reward probability (1 and 0.2 respectively for 100R and 20R trials and 0 otherwise). Event-related predictors were convolved with temporal basis functions, allowing each predictor to explain signal fluctuations over a defined interval following the corresponding task event. Temporal kernels were represented by seven cubic B-spline basis functions to allow flexible temporal influence of the event predictors. Each temporally expanded predictor was thus constructed as follows:

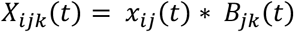

Where *x*_*ij*_(*t*) is the event vector, *B*_*jk*_ (*t*) is the *k*-th B-spline basis function for predictor *j* and * denotes convolution. We assessed the robustness of model performance to kernel duration across a range of cue- and outcome-associated kernel durations. In the final model, cue-onset and cue-associated reward probability kernels extended for 2s following cue onset, while outcome kernels extended for 4.5s following trial outcome event. No temporal expansion was applied to the lick rate predictor.

Models were fitted using ordinary least-squares regression and evaluated using five-fold cross-validation, with trials randomly assigned to folds such that all time points belonging to an individual trial remained together in either the training or test set. Predictive performance was quantified from the concatenated held-out predictions as the squared Pearson correlation (*r*^2^) between observed and predicted neural activity. Squared correlation was used as it measures the correspondence between observed and predicted neural traces without requiring a one-to-one match of their absolute amplitudes.

The contribution of each predictor was quantified by removing all design-matrix components belonging to that predictor, refitting the remaining model, and calculating the reduction in cross-validated predictive performance relative to the full model as follows:

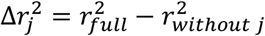

### Functional regression

To characterise the within-trial contribution of the primary predictors of interest (reward probability and licking), we additionally performed functional regression analysis focussing on only the reward-associated trials (100R & 20R). We specifically estimated the time-resolved regression coefficients by modelling the photometry activity at each time point within the trial using multiple linear regression with three predictors – reward probability, lick rate and an outcome predictor as follows:

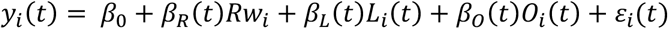

Where *y*_*i*_(*t*) is the neural signal on trial *i* at time *t, Rw*_*i*_ represents cue-associated reward probability, *L*_*i*_(*t*) is the instantaneous lick rate and *O*_*i*_(*t*) is the realised outcome. Reward probability was represented at the trial level according to cue-associated reward probability similar to that in the encoding model (1.0 for 100R trials and 0.2 for 20R trials), while reward delivery and reward omission were represented by separate outcome regressors beginning at the session-specific median outcome time. The model was fitted using ordinary least squares across trials independently at each time point. This resulted in generating a time-varying coefficient, *β*_*j*_(*t*), for each predictor *j*, that describes its relationship with neural activity over the course of the trial while accounting for other predictors. Note that in this approach, a separate cue-onset predictor was not included as condition-independent trial-locked activity at onset was absorbed by the intercept term, *β*_0_, while *β*_*R*_(*t*) captured the differences associated with reward probability.

Uncertainty in the coefficient trajectories was estimated by bootstrap resampling of trials within each recording session. Trials were sampled with replacement, stratified across 100R, rewarded 20R and unrewarded 20R trials, and the complete time-resolved regression was refitted for each bootstrap sample. We resampled the whole trial to preserve the within-trial temporal structure of the neural signal and predictors. For each predictor and time point, the resulting bootstrap distribution of coefficients was used to obtain the median coefficient and 95% confidence interval. We performed this analysis independently for each mouse and training session, allowing the temporal structure of predictor-related neural activity to be visualized both within individual sessions and across learning.

#### Cluster-based permutation analysis

To visualize changes across learning, we arranged time-resolved coefficient trajectories from individual sessions by training day to generate two-dimensional maps of regression coefficients as a function of within-trial time and training day. Bootstrap confidence intervals were additionally used to identify time periods in which coefficient estimates were reliably different from zero. To account for temporal dependence between adjacent time points, significant periods of predictor-related activity were further assessed using a cluster-based permutation analysis^42^.

For each predictor, a reduced model excluding the predictor of interest was fitted and its residual trial trajectories were randomly permuted across trials before being added back to the reduced-model predictions. The full regression model was then refitted to each permuted dataset to generate a null distribution of time-resolved test statistics. Time points at which the coefficient exceeded a two-sided threshold of *p* < 0.05 were first identified, and adjacent supra-threshold time points with the same sign were grouped into temporal clusters. The strength of each cluster was summarized by its cluster mass, calculated as the sum of the absolute *t*-statistics across all time points within the cluster. Thus, cluster mass reflected both the magnitude and temporal extent of a sustained predictor-related effect.

For each permutation, the largest cluster mass observed anywhere within the analysed trial time window was retained to generate the null distribution of maximum cluster masses. An observed cluster was considered significant when *P*_*cluster*_ < 0.05, where:

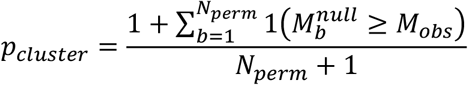

Where *M*_*obs*_ is the mass of the observed temporal cluster being assessed, 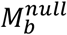 is the maximum cluster mass obtained anywhere in the analysed trial time window for *b*-th permutation, and *N*_*Perm*_ is the total number of permutations (5000 here). This finite-permutation corrected *p*-value thus tells us how frequently a cluster at least as strong as the cluster being assessed would be observed anywhere within the trial just by chance.

### Statistics

Learning-related effects were assessed using restricted permutation tests. For each hypothesis, nested linear models were constructed comprising a reduced model lacking the effect of interest and a full model containing that effect. Model improvement was quantified using a partial *F* statistic,

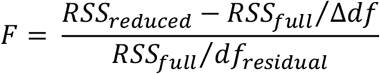

Where *RSS* denotes residual sum of squares, Δ*df* is the number of parameters added in the full model, and *df*_residual_ is its residual degrees of freedom.

We generated an empirical null distribution by restricted permutation of the data. Permutations were constrained within individual animals to preserve the repeated-measures structure of the experiment.

The models were refitted following each permutation and the partial *F* statistic recalculated. The permutation *p*-value (*p*_PERM_) was calculated as

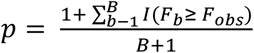

Where *B* = 5000 permutations, *F*_obs_ is the observed statistic and *F*_b_ denotes the statistic obtained from *b*^th^ permutation. Permutations were performed according to the specific null hypothesis being tested in the respective analysis.

For neural response analysis (Fig. 2), we tested three complementary hypotheses. To assess an overall reward-condition effect, we randomly exchanged the 100R and 20R condition labels within each matched mouse × training-day session, preserving the paired structure of the observations. To assess an overall training-day effect, we shuffled the categorical day labels within each mouse while keeping the paired 100R and 20R observations from each session together. This allowed us to test the effect of training-day/learning without explicitly assuming a linear trajectory. The training-day × reward-condition interaction was tested using the within-session condition contrast,

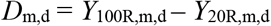

as plotted in Fig 2F. *Y* is the respective neural response metric per mouse (*m*) per day (*d*). The reduced model contained mouse identity, whereas the full model additionally contained categorical training day. We shuffled the training-day labels of the contrasts within a mouse to generate the null distribution. Thus, this test assessed whether the magnitude of the 100R-20R response difference varied across training. The 20R omission response, for which no corresponding second condition was present, was tested for a categorical training-day effect using the same within-mouse day-permutation procedure.

For assessing learning-related changes in neural-lick coupling (Fig 3), we accounted for the fact the metric is a session-wise regression coefficient, that is, an *estimated* parameter. Because these coefficients were estimated with differing precision, we performed model fitting with inverse-variance weighted coefficients derived from their standard errors. Separate permutation tests assessed whether coupling varied across categorical training day within each reward condition. Differences in learning trajectory between reward conditions were assessed by testing the Day × Condition interaction.

Finally, for assessing the learning-related changes in the predictor contribution in the encoding model analysis (Fig 4), Δr^*2*^ values were fitted with categorical training-day. Reduced models containing mouse identity were compared with full models additionally containing categorical day, and day labels were shuffled within individual mice to generate the empirical null distribution. We thus tested whether the contribution of each model component varied systematically across learning without assuming a linear change.

---

**Fig S1:**
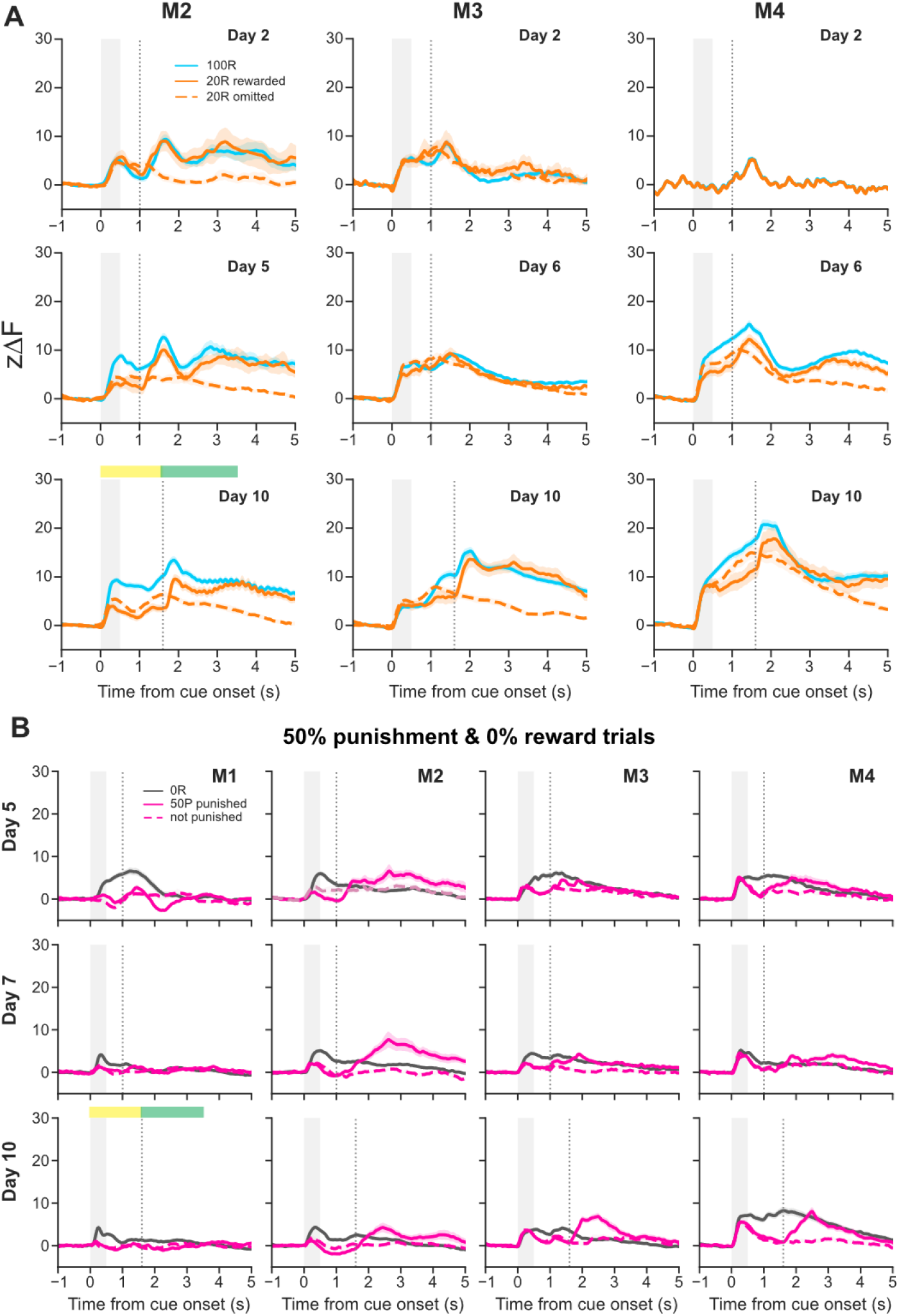
SC neural activity across training. **A)** Trial-averaged calcium photometry traces for the other mice across training. Yellow and green bars depict the pre-outcome and the outcome windows respectively used for quantifying epoch-wise response modulation metrics represented in Fig 2. All mice exhibit a diverging neural activity for the two rewarded conditions during the pre-outcome phase (yellow bar). **B)** Trial-averaged calcium photometry traces for all mice for the no-reward and the punishment trials. The *y*-axes are scaled to match the rewarded trial plots (above and Fig 2A) to aid in amplitude comparison. The neural activity exhibited very little amplitude fluctuations, though the pre-outcome responses to the punishment trials were lower than the no-reward trials. These were accounted for in the encoding model.

**Fig S2:**
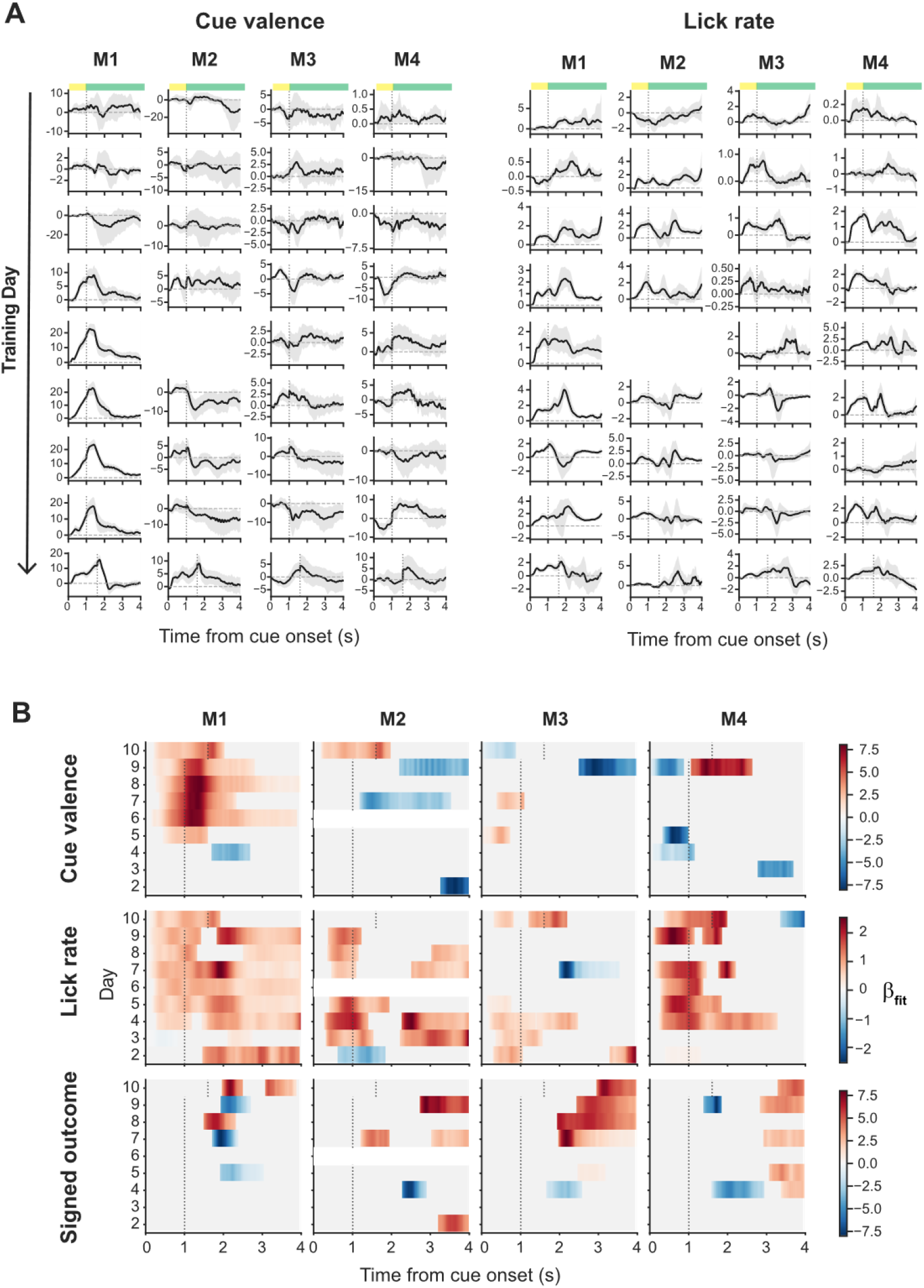
Functional regression. **A)** Time-resolved regression coefficients (*β*) for a reward probability-weighted cue predictor and a lick predictor are plotted across training day for all mice (shaded region around the curves represent 95% confidence interval). Yellow and green horizontal bars on top represent the pre-outcome and outcome epochs respectively. While the precise temporal dynamics of the trajectories exhibit heterogeneity across mice, *β* curves corresponding to the pre-outcome/anticipatory period for the both predictors show a restructuring over learning. **B)** On plotting the amplitudes of the regression coefficients when they reached significance (Methods), we see that both the coefficient values and their within-trial temporal dynamics varied across mice. Both cue valence and licking contributed to the anticipatory neural activity with increased contribution in the pre-outcome window over learning.

## REFERENCES

1. Dickinson, A., and Balleine, B. (1994). Motivational control of goal-directed action. Animal Learning & Behavior 22, 1–18. 10.3758/BF03199951.

2. Schultz, W. (2015). Neuronal Reward and Decision Signals: From Theories to Data. Physiological Reviews 95, 853–951. 10.1152/physrev.00023.2014.

3. Schultz, W., Dayan, P., and Montague, P.R. (1997). A Neural Substrate of Prediction and Reward. Science 275, 1593–1599. 10.1126/science.275.5306.1593.

4. Watabe-Uchida, M., Eshel, N., and Uchida, N. (2017). Neural Circuitry of Reward Prediction Error. Annual Review of Neuroscience 40, 373–394. 10.1146/annurev-neuro-072116-031109.

5. Cox, J., and Witten, I.B. (2019). Striatal circuits for reward learning and decision-making. Nat Rev Neurosci 20, 482–494. 10.1038/s41583-019-0189-2.

6. Drieu, C., Zhu, Z., Wang, Z., Fuller, K., Wang, A., Elnozahy, S., and Kuchibhotla, K. (2025). Rapid emergence of latent knowledge in the sensory cortex drives learning. Nature 641, 960–970. 10.1038/s41586-025-08730-8.

7. Wang, A.Y., Miura, K., and Uchida, N. (2013). The dorsomedial striatum encodes net expected return, critical for energizing performance vigor. Nat Neurosci 16, 639–647. 10.1038/nn.3377.

8. Ramakrishnan, A., Byun, Y.W., Rand, K., Pedersen, C.E., Lebedev, M.A., and Nicolelis, M.A.L. (2017). Cortical neurons multiplex reward-related signals along with sensory and motor information. Proceedings of the National Academy of Sciences 114, E4841–E4850. 10.1073/pnas.1703668114.

9. Basso, M.A., and May, P.J. (2017). Circuits for Action and Cognition: A View from the Superior Colliculus. Annual Review of Vision Science 3, 197–226. 10.1146/annurev-vision-102016-061234.

10. Masullo, L., Mariotti, L., Alexandre, N., Freire-Pritchett, P., Boulanger, J., and Tripodi, M. (2019). Genetically Defined Functional Modules for Spatial Orienting in the Mouse Superior Colliculus. Current Biology 29, 2892–2904.e8. 10.1016/j.cub.2019.07.083.

11. Wilson, J.J., Alexandre, N., Trentin, C., and Tripodi, M. (2018). Three-Dimensional Representation of Motor Space in the Mouse Superior Colliculus. Current Biology 28, 1744–1755.e12. 10.1016/j.cub.2018.04.021.

12. González-Rueda, A., Jensen, K., Noormandipour, M., de Malmazet, D., Wilson, J., Ciabatti, E., Kim, J., Williams, E., Poort, J., Hennequin, G., et al. (2024). Kinetic features dictate sensorimotor alignment in the superior colliculus. Nature 631, 378–385. 10.1038/s41586-024-07619-2.

13. Griggs, W.S., Amita, H., Gopal, A., and Hikosaka, O. (2018). Visual Neurons in the Superior Colliculus Discriminate Many Objects by Their Historical Values. Front. Neurosci. 12. 10.3389/fnins.2018.00396.

14. Ikeda, T., and Hikosaka, O. (2003). Reward-Dependent Gain and Bias of Visual Responses in Primate Superior Colliculus. Neuron 39, 693–700. 10.1016/S0896-6273(03)00464-1.

15. Ikeda, T., and Hikosaka, O. (2007). Positive and Negative Modulation of Motor Response in Primate Superior Colliculus by Reward Expectation. Journal of Neurophysiology 98, 3163–3170. 10.1152/jn.00975.2007.

16. Baruchin, L.J., Alleman, M., and Schröder, S. (2023). Reward Modulates Visual Responses in the Superficial Superior Colliculus of Mice. J. Neurosci. 43, 8663–8680. 10.1523/JNEUROSCI.0089-23.2023.

17. Comoli, E., Coizet, V., Boyes, J., Bolam, J.P., Canteras, N.S., Quirk, R.H., Overton, P.G., and Redgrave, P. (2003). A direct projection from superior colliculus to substantia nigra for detecting salient visual events. Nat Neurosci 6, 974–980. 10.1038/nn1113.

18. Takakuwa, N., Kato, R., Redgrave, P., and Isa, T. (2017). Emergence of visually-evoked reward expectation signals in dopamine neurons via the superior colliculus in V1 lesioned monkeys. eLife 6, e24459. 10.7554/eLife.24459.

19. Zhang, Y.-F., Dufour, J.-P., Zatka-Haas, P., Redgrave, P., Black, M.J., Lak, A., Mann, E., Cragg, S.J., Abraham, W.C., and Reynolds, J.N. (2026). The superior colliculus gates dopamine responses to conditioned stimuli in visual classical conditioning. Nat Commun 17, 5885. 10.1038/s41467-026-72167-4.

20. Poisson, C.L., Green, I.K., Stemmler, G.M., Prohofsky, J., Wolff, A.R., Herubin, C., Blake, M., and Saunders, B.T. (2025). Superior Colliculus Projections Drive Dopamine Neuron Activity and Movement But Not Value. J. Neurosci. 45. 10.1523/JNEUROSCI.0291-25.2025.

21. Rescorla, R.A., and Wagner, A.R. (1972). A theory of Pavlovian conditioning: Variations in the effectiveness of reinforcement and nonreinforcement. In Classical Conditioning II (A.H. Black & W.F. Prokasy, Eds.), pp. 64–99.

22. Fiorillo, C.D., Tobler, P.N., and Schultz, W. (2003). Discrete Coding of Reward Probability and Uncertainty by Dopamine Neurons. Science 299, 1898–1902. 10.1126/science.1077349.

23. Steinberg, E.E., Keiflin, R., Boivin, J.R., Witten, I.B., Deisseroth, K., and Janak, P.H. (2013). A causal link between prediction errors, dopamine neurons and learning. Nat Neurosci 16, 966–973. 10.1038/nn.3413.

24. Weldon, D.A., Patterson, C.A., Colligan, E.A., Nemeth, C.L., and Rizio, A.A. (2008). Single unit activity in the rat superior colliculus during reward magnitude task performance. Behavioral Neuroscience 122, 183–190. 10.1037/0735-7044.122.1.183.

25. Weldon, D.A., DiNieri, J.A., Silver, M.R., Thomas, A.A., and Wright, R.E. (2007). Reward-related neuronal activity in the rat superior colliculus. Behavioural Brain Research 177, 160–164. 10.1016/j.bbr.2006.11.004.

26. Lee, J., and Sabatini, B.L. (2021). Striatal indirect pathway mediates exploration via collicular competition. Nature 599, 645–649. 10.1038/s41586-021-04055-4.

27. Rossi, M.A., Li, H.E., Lu, D., Kim, I.H., Bartholomew, R.A., Gaidis, E., Barter, J.W., Kim, N., Cai, M.T., Soderling, S.H., et al. (2016). A GABAergic nigrotectal pathway for coordination of drinking behavior. Nat Neurosci 19, 742–748. 10.1038/nn.4285.

28. Ito, B.S., Gao, Y., Kardon, B., and Goldberg, J.H. (2025). A collicular map for touch-guided tongue control. Nature 637, 1143–1151. 10.1038/s41586-024-08339-3.

29. Gopal, A., and Hikosaka, O. (2026). Multiple groups of neurons in the superior colliculus convert value signals into saccadic vigor. iScience 29, 114563. 10.1016/j.isci.2025.114563.

30. Benavidez, N.L., Bienkowski, M.S., Zhu, M., Garcia, L.H., Fayzullina, M., Gao, L., Bowman, I., Gou, L., Khanjani, N., Cotter, K.R., et al. (2021). Organization of the inputs and outputs of the mouse superior colliculus. Nat Commun 12, 4004. 10.1038/s41467-021-24241-2.

31. Horwitz, G.D., and Newsome, W.T. (1999). Separate Signals for Target Selection and Movement Specification in the Superior Colliculus. Science 284, 1158–1161. 10.1126/science.284.5417.1158.

32. Carello, C.D., and Krauzlis, R.J. (2004). Manipulating Intent: Evidence for a Causal Role of the Superior Colliculus in Target Selection. Neuron 43, 575–583. 10.1016/j.neuron.2004.07.026.

33. Wang, L., McAlonan, K., Goldstein, S., Gerfen, C.R., and Krauzlis, R.J. (2020). A Causal Role for Mouse Superior Colliculus in Visual Perceptual Decision-Making. J. Neurosci. 40, 3768–3782. 10.1523/JNEUROSCI.2642-19.2020.

34. Felsen, G., and Mainen, Z.F. (2008). Neural Substrates of Sensory-Guided Locomotor Decisions in the Rat Superior Colliculus. Neuron 60, 137–148. 10.1016/j.neuron.2008.09.019.

35. Duan, C.A., Pagan, M., Piet, A.T., Kopec, C.D., Akrami, A., Riordan, A.J., Erlich, J.C., and Brody, C.D. (2021). Collicular circuits for flexible sensorimotor routing. Nat Neurosci 24, 1110–1120. 10.1038/s41593-021-00865-x.

36. Campagner, D., Vale, R., Tan, Y.L., Iordanidou, P., Pavón Arocas, O., Claudi, F., Stempel, A.V., Keshavarzi, S., Petersen, R.S., Margrie, T.W., et al. (2023). A cortico-collicular circuit for orienting to shelter during escape. Nature 613, 111–119. 10.1038/s41586-022-05553-9.

37. Shang, C., Liu, A., Li, D., Xie, Z., Chen, Z., Huang, M., Li, Y., Wang, Y., Shen, W.L., and Cao, P. (2019). A subcortical excitatory circuit for sensory-triggered predatory hunting in mice. Nat Neurosci 22, 909–920. 10.1038/s41593-019-0405-4.

38. Kim, J., Beltramo, R., and Poort, J. (2026). Contributions of superior colliculus and primary visual cortex to visual spatial detection in freely moving mice. Current Biology 36, 1562–1570.e3. 10.1016/j.cub.2026.01.077.

39. Solié, C., Contestabile, A., Espinosa, P., Musardo, S., Bariselli, S., Huber, C., Carleton, A., and Bellone, C. (2022). Superior Colliculus to VTA pathway controls orienting response and influences social interaction in mice. Nat Commun 13, 817. 10.1038/s41467-022-28512-4.

40. Gershman, S.J., and Uchida, N. (2019). Believing in dopamine. Nat Rev Neurosci 20, 703–714. 10.1038/s41583-019-0220-7.

41. Mathis, A., Mamidanna, P., Cury, K.M., Abe, T., Murthy, V.N., Mathis, M.W., and Bethge, M. (2018). DeepLabCut: markerless pose estimation of user-defined body parts with deep learning. Nat Neurosci 21, 1281–1289. 10.1038/s41593-018-0209-y.

42. Maris, E., and Oostenveld, R. (2007). Nonparametric statistical testing of EEG- and MEG-data. Journal of Neuroscience Methods 164, 177–190. 10.1016/j.jneumeth.2007.03.024.

